# Infection of the maternal-fetal interface and vertical transmission following low-dose inoculation of pregnant rhesus macaques (*Macaca mulatta*) with an African-lineage Zika virus

**DOI:** 10.1101/2022.05.16.492053

**Authors:** Michelle R. Koenig, Ann M. Mitzey, Terry K. Morgan, Xiankun Zeng, Heather A. Simmons, Andres Mejia, Fernanda Leyva Jaimes, Logan T. Keding, Chelsea M. Crooks, Andrea M. Weiler, Ellie K. Bohm, Matthew T. Aliota, Thomas C. Friedrich, Emma L. Mohr, Thaddeus G. Golos

**Affiliations:** Department of Comparative Biosciences, University of Wisconsin-Madison, Madison, Wisconsin, United States of America; Department of Pathology, Oregon Health and Science University, Portland, Oregon, United States of America; Department of Obstetrics and Gynecology, Oregon Health and Science University, Portland, Oregon, United States of America; Pathology Division, United States Army Medical Research Institute of Infectious Diseases, Frederick, MD, United States of America; Wisconsin National Primate Research Center, University of Wisconsin-Madison, Madison, WI, United States of America; Department of Obstetrics and Gynecology, University of Wisconsin-Madison, Madison, Wisconsin, United States of America; Department of Pathobiological Sciences, University of Wisconsin-Madison, Madison, Wisconsin, United States of America; Department of Veterinary and Biomedical Sciences, University of Minnesota, Twin Cities, St. Paul, Minnesota, United States of America; Department of Pediatrics, University of Wisconsin-Madison, Madison, Wisconsin, United States of America

## Abstract

**Background:** Congenital Zika virus (ZIKV) infection can result in birth defects, including malformations in the fetal brain and visual system. There are two distinct genetic lineages of ZIKV: African and Asian. Asian lineage ZIKVs have been associated with adverse pregnancy outcomes in humans; however, recent evidence from experimental models suggests that African-lineage viruses can also be vertically transmitted and cause fetal harm.

**Methodology/Principal Findings:** To evaluate the potential for vertical transmission of African-lineage ZIKV, we inoculated nine pregnant rhesus macaques (*Macaca mulatta*) subcutaneously with 44 plaque-forming units of a ZIKV strain from Senegal, (ZIKV-DAK). Dams were inoculated either at gestational day 30 or 45. Following maternal inoculation, pregnancies were surgically terminated seven or 14 days later and fetal and maternal-fetal interface tissues were collected and evaluated. Infection in the dams was evaluated via plasma viremia and neutralizing antibody titers pre- and post- ZIKV inoculation. All dams became productively infected and developed strong neutralizing antibody responses. ZIKV RNA was detected in maternal-fetal interface tissues (placenta, decidua, and fetal membranes) by RT-qPCR and *in situ* hybridization. *In situ* hybridization detected ZIKV predominantly in the decidua and revealed that the fetal membranes may play a role in ZIKV vertical transmission. Infectious ZIKV was detected in the amniotic fluid of three pregnancies and one fetus had ZIKV RNA detected in multiple tissues. No significant pathology was observed in any fetus; however, we did find an increase in the occurrence of decidual vasculitis and necrosis in ZIKV-exposed pregnancies compared to gestational-age-matched controls.

**Conclusions/Significance:** This study demonstrates that African-lineage ZIKV, like Asian-lineage ZIKV, can be vertically transmitted to the macaque fetus during pregnancy. The low inoculating dose used in this study suggests a low minimal infectious dose for rhesus macaques. Vertical transmission with a low dose in macaques further supports the high epidemic potential of African ZIKV strains.

**Author Summary:** Zika virus infection during pregnancy can result in adverse pregnancy outcomes including birth defects and miscarriage. There are two distinct genetic backgrounds of Zika virus: Asian-lineage and African-lineage. Currently, only Asian-lineage Zika virus is causally associated with adverse pregnancy outcomes in people. However, experimental studies have shown that African-lineage Zika virus can infect the fetus during pregnancy and cause adverse outcomes. Adverse pregnancy outcomes may not be associated with infection in people until there is an outbreak in a naive population. Thus, as African-lineage Zika virus continues to spread globally, the risk that it may pose to pregnant people remains a public health concern. In this study, we demonstrate that African-lineage Zika virus can be transmitted from the mother to the fetus during pregnancy. This study is significant because we used rhesus macaques, an animal that shares many key elements of Zika virus infection in pregnant people. This study is also significant because we inoculated pregnant macaques with a very small amount of virus, suggesting that fetal infection reported in previously published macaque studies is not limited to high-dose inoculation.

## Introduction

Zika virus (ZIKV) is a flavivirus that is primarily transmitted by *Aedes spp.* mosquitoes [1]. ZIKV was initially discovered in Uganda in 1947 in a sentinel rhesus macaque (*Macaca mulatta*) [2]. ZIKV has since spread from the African continent to Asia, the Pacific, and the Americas [3]. Two distinct genetic lineages of ZIKV have been identified, African-lineage and Asian-lineage. Asian-lineage viruses spread from Asia to the Pacific where it was detected in 2007 [3] and was introduced to Brazil between 2013 and 2014 [4]. The introduction of ZIKV in Brazil led to an outbreak in 2015 [4]. During this outbreak, ZIKV was first causally connected to birth defects and other adverse pregnancy outcomes [5].

In utero ZIKV exposure is associated with malformations in the fetal brain and visual system, along with fetal demise and pregnancy loss [6], [7], [8], [9], [10], [11], [12]. Of the two distinct genetic lineages of ZIKV, only Asian-lineage ZIKV is currently associated with congenital infection in humans [13]. Recently, African-lineage ZIKV has been detected in Brazil [14][15] [16]. The novel detection of African-lineage ZIKV outside the African continent demonstrates the potential of a new outbreak and underscores the need to further evaluate the threat that African-lineage ZIKV poses to pregnant people.

Congenital birth defects or adverse pregnancy outcomes have not been formally associated with African-lineage ZIKV infection during pregnancy in people. However, *in vivo* studies done in mice and macaques suggest that African-lineage ZIKV could pose a substantial risk when infection occurs during pregnancy [17] [18] [19] [20]. African-lineage infection in pregnant immune-deficient mice revealed the potential for vertical transmission during pregnancy [17][18]. In contrast to Asian-lineage ZIKV, African-lineage ZIKV infection in pregnant mice is more likely to cause fetal demise than birth defects [17] [18]. Aubry et al. reported higher rates of fetal reabsorption in mouse pregnancies infected with African-lineage ZIKV, whereas infection with Asian-lineage resulted in varying capacities to cause fetal harm including fetal brain abnormalities as well as fetal death/resorption [17] [21].

The rhesus macaque model accurately reflects many key aspects of ZIKV infection during human pregnancy, thus making it a highly translatable model. Previous macaque studies have offered unique insights into the potential of African ZIKV infection to cause fetal harm during pregnancy [19] [20] [22]. These studies have all used the same low-passage African-lineage ZIKV strain from Senegal (ZIKV-DAK) yet have used a range of inoculation doses and routes, and gestational timing of infection. Crooks et al. found that a subcutaneous maternal inoculation of 1X10^4^ plaque forming units (PFU) at gestation day (gd) 45 resulted in a higher viral burden in the placenta than the same dose of an Asian-lineage virus [19], but did not find any evidence of vertical transmission. Newman et al. showed that maternal vaginal inoculation with 3X10^8^ PFU at gd 30 resulted in vertical transmission and fetal loss in two of three pregnancies [20]. Raasch et al. demonstrated that a maternal subcutaneous dose of 1X10^8^ gd 45 resulted in vertical transmission and fetal demise in three out of eight pregnancies [22]. The Crooks et al. and Raasch et al. studies suggest that vertical transmission may be dose-dependent. However, these studies evaluated the fetuses/neonates and maternal-fetal interface (MFI) tissues either when fetal death was detected or at gestational term, more than 15 weeks after maternal ZIKV inoculation.

Collectively, these data underscore the potential of African-lineage ZIKV to adversely affect the fetus in utero. To further evaluate the capacity of African-lineage ZIKV to infect maternal-fetal interface tissues and cause fetal harm, we inoculated nine pregnant rhesus macaques with a relatively low dose of 44 PFU of ZIKV-DAK and terminated the pregnancies within two weeks post-inoculation. This design allows us to capture the early stages of infection in the MFI and vertical transmission and evaluate their vulnerability to African-lineage ZIKV. Furthermore, assessing the fetuses within two weeks of maternal ZIKV inoculation allows us to investigate viral burden and pathology in the fetuses before fetal loss is likely to occur. We found that this low dose of African-lineage ZIKV is capable of infecting macaques and causing vertical transmission. This study indicates a low minimal infectious dose for African-lineage ZIKV in rhesus macaques and suggests a high epidemic potential of African ZIKV strains.

## Materials and Methods

### Experimental Design

A total of nine pregnant female rhesus macaques (*Macaca mulatta*) were subcutaneously inoculated with 44 PFU of a Senegal isolate of African-lineage Zika virus ZIKV/*Aedes africanus*/SEN/DAK-AR-41524/1984 (ZIKV-DAK), Genbank accession number KY348860, during early pregnancy. Five dams were infected at approximately gd 30, two of these pregnancies were terminated seven days post-infection (dpi), and three were terminated 14 dpi. Four dams were infected later in pregnancy, at approximately gd 45 followed by pregnancy termination at 14 dpi. A total of seven pregnancies terminated between gd 37 and gd 61 were used for controls for histological evaluation. Dams with control pregnancies were inoculated with saline and subjected to the same experimental regimen as the ZIKV infected pregnancies. Details on the timing of ZIKV/saline inoculation and pregnancy termination are provided in S1 Table.

### Ethics

The rhesus macaques used in this study were cared for by the staff at the Wisconsin National Primate Research Center (WNPRC) according to regulations and guidelines of the University of Wisconsin Institutional Animal Care and Use Committee, which approved this study protocol (G005691) in accordance with recommendations of the Weatherall report and according to the principles described in the National Research Council’s Guide for the Care and Use of Laboratory Animals. All animals were housed in enclosures with at least 4.3, 6.0, or 8.0 sq. ft. of floor space, measuring 30, 32, or 36 inches high, and containing a tubular PVC or stainless steel perch. Each individual enclosure was equipped with a horizontal or vertical sliding door, an automatic water lixit, and a stainless steel feed hopper. All animals were fed using a nutritional plan based on recommendations published by the National Research Council. Twice daily, macaques were fed a fixed formula, extruded dry diet (2050 Teklad Global 20% Protein Primate Diet) with adequate carbohydrate, energy, fat, fiber (10%), mineral, protein, and vitamin content. Dry diets were supplemented with fruits, vegetables, and other edible foods (e.g., nuts, cereals, seed mixtures, yogurt, peanut butter, popcorn, marshmallows, etc.) to provide variety to the diet and to inspire species-specific behaviors such as foraging. To further promote psychological well-being, animals were provided with food enrichment, human-to-monkey interaction, structural enrichment, and manipulanda. Environmental enrichment objects were selected to minimize chances of pathogen transmission from one animal to another and from animals to care staff. While on study, all animals were evaluated by trained animal care staff at least twice daily for signs of pain, distress, and illness by observing appetite, stool quality, activity level, and physical condition. Animals exhibiting abnormal presentation for any of these clinical parameters were provided appropriate care by attending veterinarians.

### Care & Use of Macaques

The female macaques described in this report were co-housed with a compatible male and observed daily for menses and breeding. Pregnancy was detected by abdominal ultrasound, and gestational age was estimated as previously described [23]. For physical examinations, virus inoculations, and blood collections, dams were anesthetized with an intramuscular dose of ketamine (10 mg/kg). Blood samples from the femoral or saphenous vein were obtained using a vacutainer system or needle and syringe. Pregnant macaques were monitored daily prior to and after inoculation for any clinical signs of infection (e.g., diarrhea, inappetence, inactivity, and atypical behaviors) and general well-being.

### Inoculation and monitoring

Nine pregnant dams were inoculated subcutaneously with 44 PFU of ZIKV/*Aedes africanus*/SEN/DAK-AR-41524/1984 (ZIKV-DAK). This virus was originally isolated from *Aedes africanus* mosquitoes with a round of amplification on *Aedes pseudocutellaris* cells, followed by amplification on C6/36 cells and two rounds of amplification on Vero cells. ZIKV-DAK was obtained from BEI Resources (Manassas, VA). The stock of ZIKV-DAK was evaluated using plaque assay on Vero cells as previously described [19]. Inocula were prepared from the viral stock described above. The stock was thawed, diluted in sterile PBS to 44 PFU/mL, and loaded into a 1-mL syringe that was kept on ice until inoculation. Animals were anesthetized as described above, and 1 mL of the inoculum was delivered subcutaneously over the cranial dorsum. Animals were monitored closely following inoculation for any signs of an adverse reaction. The 44- PFU dose was administered due to an unintentional dilution error. Seven pregnant dams served as controls and were inoculated subcutaneously with 1mL of sterile PBS, using the same procedure as described above. All control animals were subjected to the same sedation and blood collection schedule as the ZIKV inoculated animals.

### Pregnancy termination and tissue collection

A total of 14 dams had their pregnancies surgically terminated at gd 36–63 via laparotomy. This procedure was terminal for three dams, 45/14-4, 30/14-1 and 30/14/- C1, and thus plasma viremia and PRNT data points are not provided for 45/14-4 and 30/14-1 beyond 14 dpi. During the laparotomy procedure, the entire conceptus (fetus, placenta, fetal membranes, umbilical cord, and amniotic fluid) was removed. The tissues for all animals were dissected using sterile instruments which were changed between each organ/tissue to minimize possible cross-contamination. Each organ/tissue was evaluated grossly *in situ*, removed with sterile instruments, placed in a sterile culture dish, and dissected for histology, viral burden assay, or banked for future assays. Samples of the MFI included full-thickness center-cut sections of the primary and secondary placental disc containing decidua basalis and chorionic plate. Biopsy samples were collected for RT-qPCR evaluation including: decidua basalis dissected from the maternal surface of the placenta, placental biopsies from each placental disc with the decidua basalis removed, chorionic plate, fetal membranes, and placental bed (the uterine placental attachment site containing deep decidua basalis and uterine myometrium). Although placenta biopsies were taken from the placenta after the decidua basalis was peeled off, it is likely that a small portion of the decidua may have been still attached as well as the trophoblastic shell and portions of the chorionic plate. Half of the fetus was preserved for histological evaluation and samples from the following tissues and fluids were collected from the other half: cerebral spinal fluid (CSF), fetal brain with or without skull, eye, heart or fetal chest, fetal limb containing muscle and skin, liver, kidney, and spleen.

### Viral RNA isolation from blood, tissues, and other fluids

RNA was isolated from maternal and fetal plasma, CSF, and amniotic fluid using the Viral Total Nucleic Acid Purification Kit (Promega, Madison, WI) on a Maxwell 48 RSC instrument (Promega, Madison, WI) as previously reported [24]. Fetal and maternal tissues were processed with RNAlater (Invitrogen, Carlsbad, CA) according to the manufacturer’s protocols. RNA was recovered from tissue samples using a modification of a previously described method [25]. Briefly, up to 200 mg of tissue was disrupted in TRIzol (Life Technologies, Carlsbad, CA) with 2 x 5 mm stainless steel beads using a TissueLyser (Qiagen, Germantown, MD) for 3 minutes at 25 r/s for 2 cycles. Following homogenization, samples in TRIzol were separated using Bromo-chloro-propanol (Sigma-Aldrich, St. Louis, MO). The aqueous phase was collected, and glycogen was added as a carrier. The samples were washed in isopropanol and ethanol precipitated. RNA was re-suspended in 5 mM Tris pH 8.0 and stored at -80 °C.

### Quantitative reverse transcriptase (RT-qPCR)

ZIKV RNA was isolated from both fluid and tissue samples as previously described [18]. Viral RNA was then quantified using a highly sensitive RT-qPCR assay based on the one developed by Lanciotti et al. [26], though the primers were modified with degenerate bases at select sites to accommodate African-lineage Zika viruses.

RNA was reverse-transcribed and amplified using the TaqMan Fast Virus 1-Step Master Mix RT-qPCR kit (LifeTechnologies, Carlsbad, CA) on the LightCycler 480 or LC96 instrument (Roche, Indianapolis, IN), and quantified by interpolation onto a standard curve made up of serial tenfold dilutions of *in vitro* transcribed RNA. RNA for this standard curve was transcribed from a plasmid containing an 800 bp region of the Zika virus genome that is targeted by the RT-qPCR assay. The final reaction mixtures contained 150 ng random primers (Promega, Madison, WI), 600 nM each primer and 100 nM probe. Primer and probe sequences are as follows: forward primer: 5’- CGYTGCCCAACACAAGG-3’, reverse primer: 5′-CCACYAAYGTTCTTTTGCABACAT-3′ and probe: 5′-6-carboxyfluorescein-AGCCTACCTTGAYAAGCARTCAGACACYCAA. The limit of detection for fluids (plasma, amniotic fluid, CSF) with this assay is 100 copies/ml.

### Plaque Reduction Neutralization test (PRNT)

Macaque serum samples were assessed for ZIKV neutralizing antibodies utilizing a plaque reduction neutralization test (PRNT). Endpoint titrations of reactive sera, utilizing a 90% cutoff (PRNT90), were performed as previously described [27] against ZIKV/Aedes africanus/SEN/DAK-AR-41524/1984 (ZIKV-DAK). Briefly, ZIKV was mixed with serial 2-fold dilutions of serum for 1 hour at 37°C before being added to Vero cells, and neutralization curves were generated using GraphPad Prism software (La Jolla, CA). The resulting data were analyzed by nonlinear regression to estimate the dilution of serum required to inhibit both 90% and 50% of infection.

### Viral quantification by plaque assay

Amniotic fluid samples from 30/14-1, 30/7-2, 45/14-4, and 30/14-3 were evaluated for infectious ZIKV by plaque assay. These samples were selected based on RT-qPCR results showing a high level of ZIKV RNA. Titrations for replication competent virus quantification of amniotic fluid was completed by plaque assay on Vero cell cultures as described previously [28]. Vero cells were obtained from the American Type Culture Collection (CCL-81). Duplicate wells were infected with 0.1 mL of aliquots from serial 10-fold dilutions in growth media and virus was adsorbed for 1h. Following incubation, the inoculum was removed, and cell monolayers were overlaid with 3 mL containing a 1:1 mixture of 1.2% oxoid agar and DMEM (Gibco, Carlsbad, CA, USA) with 10% (vol/vol) FBS and 2% (vol/vol) penicillin/streptomycin. Cells were incubated at 37°C in 5% CO_2_ for 2 days for plaque development. Cell monolayers were then stained with 3 mL of overlay containing a 1:1 mixture of 1.2% oxoid agar and DMEM with 2% (vol/vol) FBS, 2% (vol/vol) penicillin/streptomycin and 0.33% neutral red (Gibco). Cells were incubated overnight at 37°C and plaques were counted. Limit of detection for the plaque assay is 0.7 log10 PFU/mL.

### Histology

To assess general pathology, tissues were fixed in 4% PFA, routinely processed, and embedded in paraffin. Paraffin sections (5 µm) were stained with hematoxylin and eosin (H&E) or used for *in situ* hybridization (ISH). Histological evaluation of fetal tissue was done by a board-certified veterinary pathologist (HAS, AM) blinded to the ZIKV RNA results. Histological evaluation of the placenta center cuts for chronic deciduitis, villitis, maternal decidual vasculitis, decidual necrosis, acute villous infarctions, and villous calcifications was done by a board-certified pathologist who specializes in human placental pathology (TKM); this pathologist was only informed of the gestational age and remained blinded to all ZIKV RNA results and pregnancy treatment.

Photomicrographs were taken using brightfield on the Nikon eclipse Ti2 (Nikon Instruments Inc., Melville, NY, U.S.A) using NIS-Elements AR software version 5.02.006 (Nikon Instruments Inc., Melville, NY, U.S.A). Scale bars were added using NIS-Elements AR and photomicrographs were white balanced using Adobe Photoshop 2020 version 21.10 (Adobe Inc, San Jose, CA, U.S.A.).

### Detention of ZIKV RNA using *in situ* hybridization (ISH)

ISH was conducted as previously described [23]. The ISH probes against the Zika virus genome were purchased commercially (Advanced Cell Diagnostics, Cat No. 468361, Newark, California, USA).

### Detection of ZIKV replication using multiplex fluorescence *in situ* hybridization (mFISH)

Multiplex fluorescence *in situ* hybridization (mFISH) was performed using the RNAscope® Fluorescent Multiplex Kit (Advanced Cell Diagnostics, Newark, CA) according to the manufacturer’s instructions with modifications. Twenty ZZ probe pairs with C1 channel (green, Cat# 463781) targeting ZIKV positive sense RNA and forty ZZ probe pairs with C3 channel (red, Cat# 467911) targeting ZIKV negative sense RNA are synthesized by Advanced Cell Diagnostics. Formalin-fixed paraffin-embedded (FFPE) tissue sections were deparaffinized using xylene and hydrated through a series of ethanol washes. These tissue sections were further heated in the antigen retrieval buffer and digested by proteinase. Sections were exposed to ISH target probes and incubated at 40°C in a hybridization oven for two hours. After rinsing, ISH signal is amplified using company-provided Pre-amplifier and Amplifier conjugated to fluorescent dye. Sections were counterstained with 4’, 6-diamidino-2-phenylindole (DAPI, Thermo Fisher Scientific, Waltham, MA, USA), mounted, and stored at 4°C until image analysis. FISH images were captured on an LSM 880 Confocal Microscope with Airyscan (Zeiss, Oberkochen, Germany) and processed using open-source ImageJ software (National Institutes of Health, Bethesda, MD, USA).

### Immunohistochemistry

CD56 and CD68 stained tissues sections were immunostained using the Benchmark XT autostainer (Ventana Medical Systems, Tucson, AZ, USA) using provided pre-diluted antibodies CD56 and CD68 (Ventana Medical Systems, Tucson, AZ, USA). For cytokeratin and CD163 staining immunostaining, sections were deparaffinized, incubated in 3% hydrogen peroxide for 15 min, and subjected to a heat induced antigen retrieval protocol (10 mM citric acid at 110°C for 15 min). Sections were immunohistochemically stained using a proprietary polymer-based peroxidase staining method (Biocare Medical, Concord, CA): sections were blocked 20 min in Background Sniper (Biocare Medical), and incubated in 1:100 mouse anti-cytokeratin CAM5.2 monoclonal antibody (item 452M-94,Sigma-Aldrich, St. Louis, MO) or 1:1000 anti- CD163 monoclonal antibody (Novus Biologicals, Centennial, CO. Ref NB110-40686) overnight at 4 degrees. Slides were processed for bound antibody detection by incubation in full-strength MACH2 Polymer-horseradish peroxidase conjugate (Biocare Medical) for 20 min and developed with Betazoid DAB chromogen (Biocare Medical) for 1.5 min. All washes were in 0.1 M Tris pH 7.5 with Tween-20 at room temp.

### Statistical analysis

All statistical analysis was conducted using Graphpad Prism 9 for macOS version 9.3.1 software. Unpaired Student’s t-test was used to determine significant differences in the timing of peak viremia, viral burden at peak viremia, and viral burden in MFI tissues.

## Results

### Low-dose ZIKV-DAK inoculation resulted in productive infection and neutralizing antibody responses in all dams

A total of nine pregnant rhesus macaques were subcutaneously inoculated with a Senegalese isolate of African-lineage ZIKV during early pregnancy (Fig 1), when the fetus is at greatest risk of congenital ZIKV infection and associated birth defects [7] [9] [11], [29] [30]. Five dams were infected at approximately gd 30 and four dams at approximately gd 45. Blood was collected from the dams prior to infection, during the first two weeks after inoculation, and intermittently until up to 87 dpi. All dams received a dose of 44 PFU; although this dose was not intentionally chosen, all dams became productively infected and developed neutralizing antibody responses.

**Fig 1.**
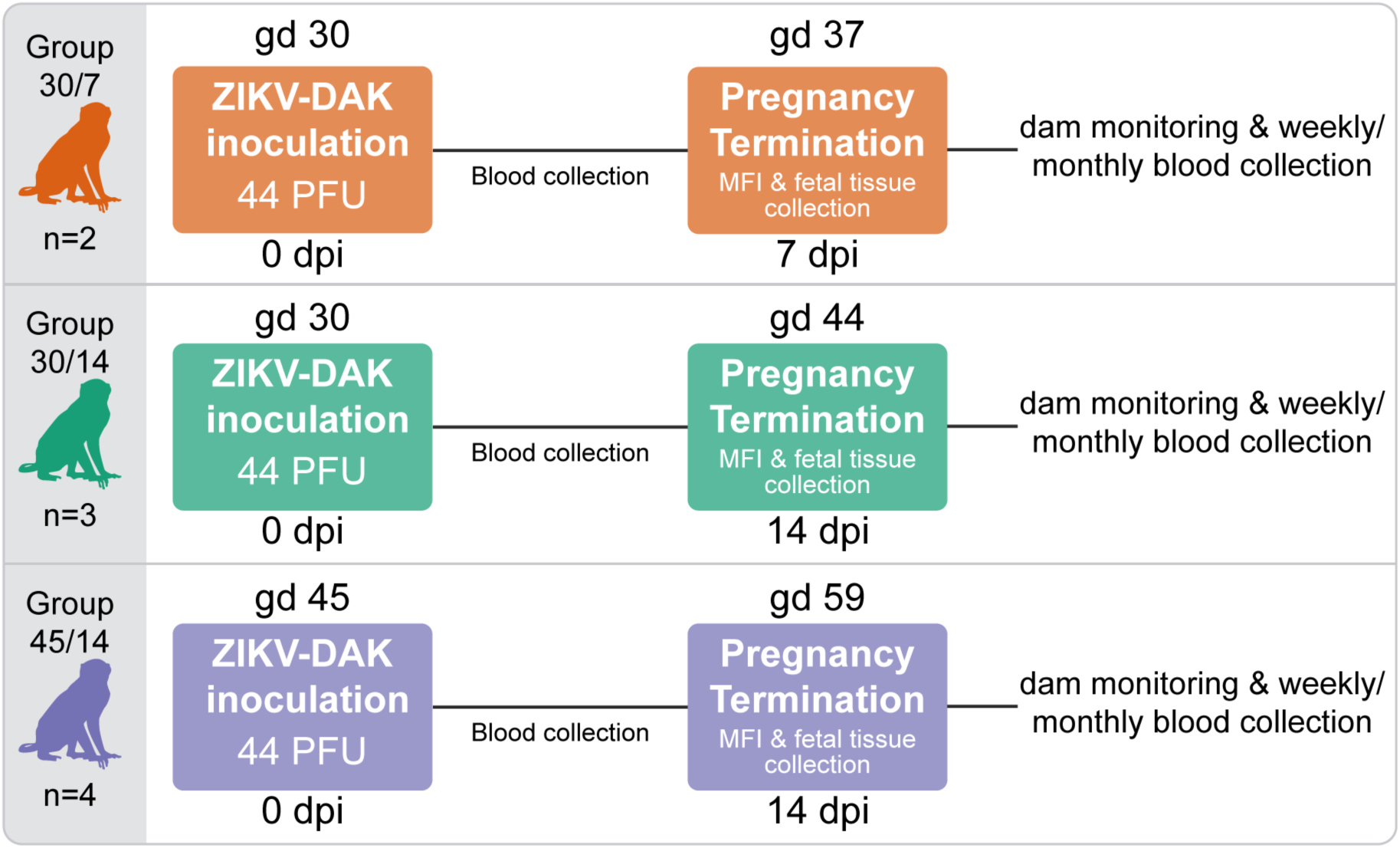
The effect of ZIKV-DAK on pregnancy was evaluated in three groups of pregnant rhesus macaques. Three groups of pregnant macaques were challenged early in gestation with 44 PFU of ZIKV-DAK at either gd 30 (Group 30/7 and Group 30/14) or gd 45 (Group 45/14). Viremia in the dams was monitored via blood collection. Pregnancies were surgically terminated at either 7 dpi (Group 30/7) or 14 dpi (Group 30/14 and Group 45/14) and MFI and fetal tissues were collected for evaluation. After pregnancy termination, the dams were monitored and blood was collected and evaluated weekly until ZIKV was no longer detected in the blood. When ZIKV was no longer detected in the blood for two consecutive blood draws, the dam was switched to monthly blood collection through 87 dpi.

All dams developed plasma viremia: seven animals resolved viremia between 10 and 16 dpi, and two dams had positive plasma viremia at 14 dpi when they were euthanized (Fig 2). Peak plasma viremia levels were between 5X10^4^ and 2.65X10^7^ ZIKV RNA copies/mL and peaked between 6 and 11 dpi. By comparing these results to a previous report that inoculated pregnant rhesus macaques with a larger dose (10^4^ PFU) of the same ZIKV isolate at gd 45 [19], we see that the animals given 44 PFU had a significant delay (p = 0.0150) in time to peak plasma viremia (S1A Fig), but reached a similar peak level of virus in the blood (S1B Fig).

**Fig 2.**
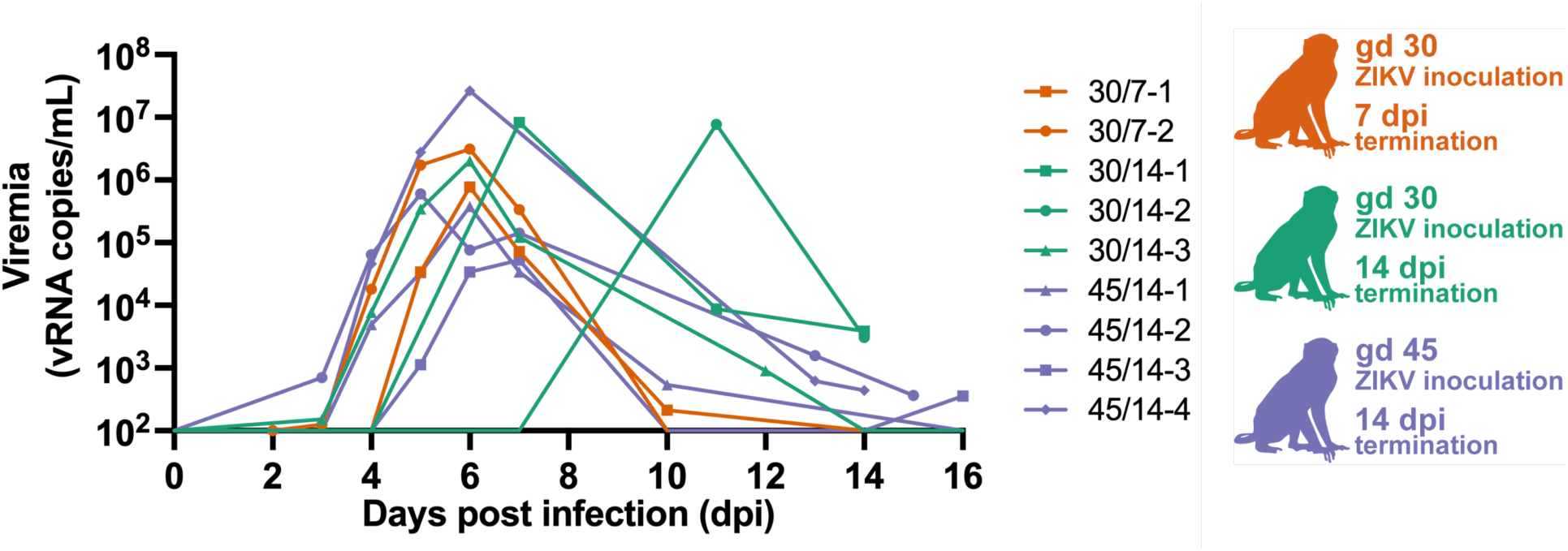
Plasma viremia following inoculation with 44 PFU of ZIKV-DAK. Plasma ZIKV loads were determined by RT-qPCR and are represented as vRNA copies/mL from 0 up to 16 dpi. Only values above the limit of detection of 100 copies/mL are shown. Experimental groups are represented by different colors, symbols represent individual animals within experimental groups.

To determine if infection with 44 PFU ZIKV was sufficient to induce neutralizing antibody (nAb) responses, we measured nAb titers in the serum prior to infection and at approximately 14 or 28 dpi. All animals had robust titers of nAb at approximately 14 or 28 dpi (Fig 3): the titers of nAb induced by inoculation with 44 PFU are not significantly different from those induced by challenge with 10^4^ PFU [19] (S3 Fig). One animal, 45/14-1, had low titers of nAb prior to infection (S2 Fig); nonetheless, this animal had a viremia profile similar to the other animals in the study (Fig 2) and its antibody responses were in the range of those produced by an inoculation dose of 10^4^ PFU (S2 Fig).

**Figure 3.**
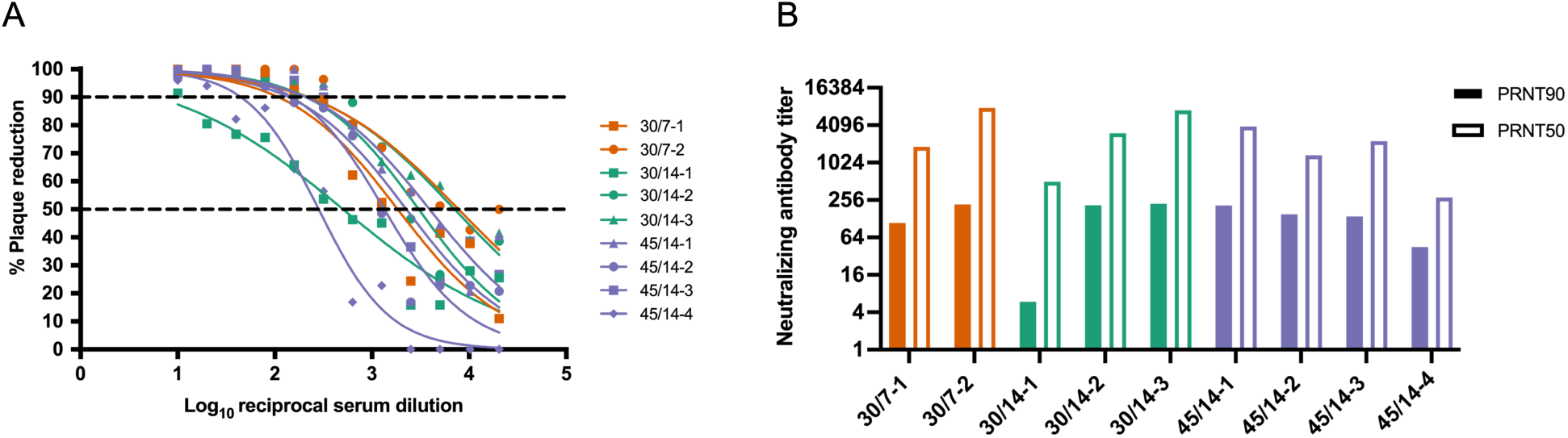
Neutralizing antibody titers following inoculation with 44 PFU of ZIKV- DAK. Plaque reduction neutralization tests (PRNT) were performed on serum samples collected at 14 dpi or approximately 28 dpi. Data are expressed relative to infectivity in the absence of serum. A) Neutralization curves. B) PRNT90 and PRNT50 values were estimated using nonlinear regression analysis and are indicated with dotted lines in Panel A.

### ZIKV-DAK was consistently detected in maternal-fetal interface tissues shortly after maternal inoculation

To assess the presence of ZIKV in the MFI we surgically terminated the pregnancies shortly after maternal inoculation. Two pregnancies were terminated at 7 dpi and seven pregnancies were terminated 14 dpi. Tissues and fluids from the MFI were evaluated for ZIKV RNA using RT-qPCR and ISH.

Collectively, ISH and RT-qPCR results show that the MFI was consistently infected with ZIKV, with virus present in the placenta, decidua basalis, and fetal membranes (Fig 4). Details on viral burden in MFI tissues are shown in S4A Fig. ZIKV burden in MFI tissues from all three experimental groups were similar (S4B Fig). ISH demonstrated that infection in the placenta was mainly confined to the trophoblastic shell and the chorionic plate. ZIKV RNA was detected within placental villi in only one pregnancy (Fig 4C). Additionally, ISH evaluation showed that the decidua basalis and trophoblastic shell were the most frequently infected MFI tissue (Fig 4C). Random sampling bias may explain discrepancies between the RT-qPCR and ISH results in the decidua basalis, chorionic plate, and fetal membranes; however, ISH evaluation suggests that infection at this stage is often focal and not widespread in the tissues.

**Fig 4.**
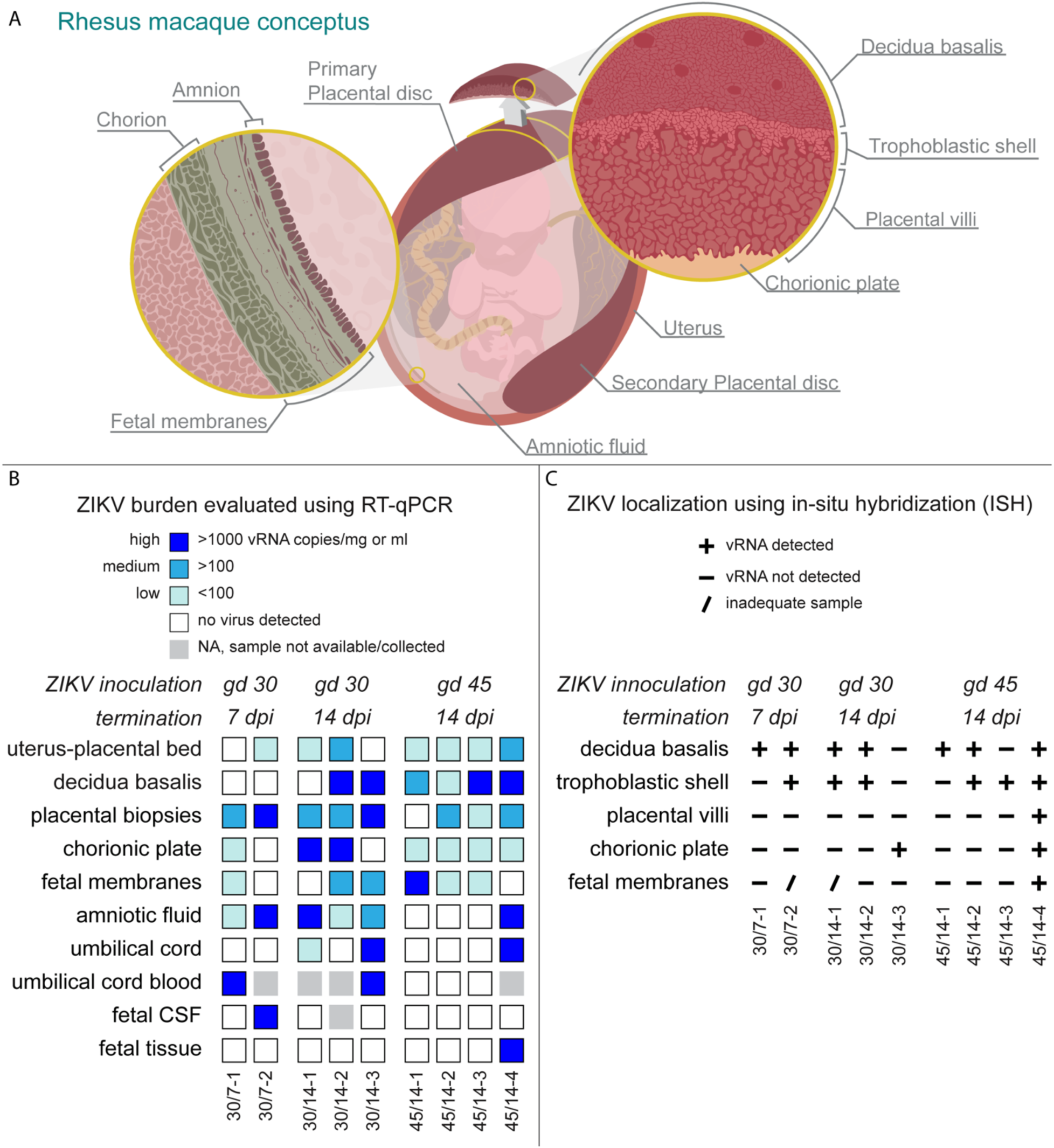
ZIKV-DAK detection in MFI and fetal tissues and fluids. A) Illustration of the rhesus macaque conceptus depicting evaluated tissues and fluids. The pregnant uterus is lined by the maternal endometrium of pregnancy called the decidua. The layers of the placenta are shown as the trophoblastic shell where the trophoblasts and the decidua meet, the placental villi, and the chorionic plate made up of the chorion and large fetal vessels. The layers of the fetal membranes are shown as the chorion and the amnion. **B)** ZIKV RNA burden as detected by RT-qPCR on RNA isolated from indicated tissues and fluids. The level of ZIKV burden is summarized as high (>1000 ZIKV RNA copies/mg of tissue or mL of fluid), medium (>100 ZIKV RNA copies/mg or mL), low (< 100 ZIKV RNA copies/mg or mL), or not detected. Placental viral loads are presented as the mean of three tissue biopsies per disc. Placental biopsies likely contained small portions of the decidua, trophoblastic shell and chorionic plate. C) ZIKV RNA presence as detected by ISH. ZIKV RNA was noted as present (+) or absent (-) in the indicated tissues. In some instances, an adequate histological sample was not obtained, thus ZIKV RNA presence or absence could not be properly evaluated (/).

### Maternal inoculation with 44 PFU of ZKV-DAK is sufficient to cause vertical transmission

In this study, we defined vertical transmission as the presence of infectious virus in the amniotic fluid. To determine the capacity of a maternal inoculation of 44 PFU to achieve vertical transmission, we evaluated the amniotic fluid with RT-qPCR and then further assessed positive samples with a plaque assay. We found infectious ZIKV in three cases three (45/14-4, 30/14-1, 30/7-2) (Fig 5A), thus confirming vertical transmission in these cases. Additionally, ZIKV RNA was detected in the umbilical cord tissue in three cases and the umbilical cord blood in two cases (Fig 4B, S4C Fig), but we were unable to confirm that the ZIKV RNA found represented a replicating virus. In the three cases with vertical transmission, ZIKV RNA was detected in multiple tissues.

**Fig 5.**
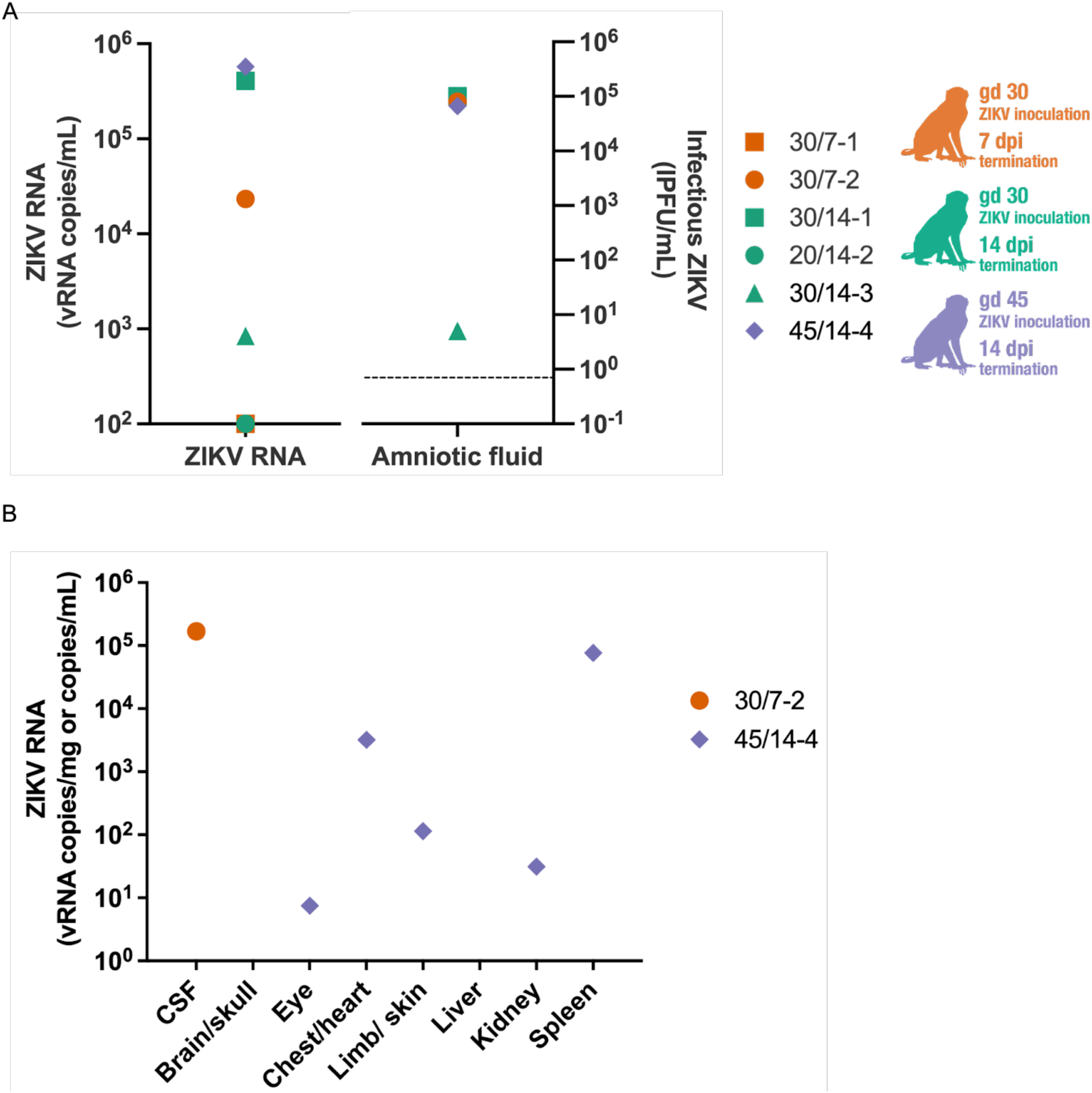
ZIKV burden in fetal tissues and fluids. A) Amniotic fluid ZIKV RNA levels determined by RT-qPCR (left panel) and infectious ZIKV determined by plaque assay (right panel). Limit of detection for the RT-qPCR is 100 copies/mL. The dashed line represents the limit of detection of 0.7 PFU/mL for the plaque assay. Four amniotic fluid samples were analyzed by plaque assay based on positive detection of vRNA by RT- qPCR. B) ZIKV RNA isolated from fetal tissues and fluids; viral loads are measured in vRNA copies/mL of fluid (CSF) or vRNA copies/mg of tissue.

In one case, 45/14-4, ZIKV RNA was detected in the CSF of the fetus in 30/7-2, and was not detected in the fetus proper in case 30/14-1 (Fig 4B, Fig 5B). We were unable to confirm that RNA detected in the fetal tissues represented replicating virus, due to the absence of negative-sense ZIKV RNA as determined using mFISH. Although no replicating virus was detected in this fetus, the presence of infectious virus in the amniotic fluid confirms that ZIKV-DAK was vertically transmitted to the fetus. Maternal inoculation at gd 30 versus gd 45 appeared to have no impact on the capacity for vertical transmission. Furthermore, pregnancies that were allowed to progress to 14 dpi compared to 7 dpi also did not demonstrate an increased likelihood of vertical transmission. Collectively, these data show that maternal inoculation of 44 PFU of ZIKV- DAK is sufficient to cause vertical transmission as early as 7 dpi.

### ZIKV-DAK may vertical transmit through the fetal membranes

To understand the pathway of vertical transmission, we used ISH to evaluate the cellular location of ZIKV RNA in histological specimens from cases that had infectious virus in the amniotic fluid (30/7-2, 30/14-1, and 45/14-4). For 30/7-2 and 30/14-1, ISH was performed on full-thickness placental center cuts taken from the primary and secondary placental discs which included decidua basalis, trophoblastic shell, placental villi, and chorionic plate with large fetal vessels. In both cases, ZIKV RNA was detected in the decidua basalis and trophoblast shell but not in the placental villi or chorionic plate (Fig 4C). Unfortunately, we were not able to obtain a high-quality histological specimen of the fetal membranes from these two cases to properly evaluate the presence of ZIKV RNA.

In the case of 45/14-4, where we have the most apparent evidence of vertical transmission to the fetus (Fig 4), ISH was performed on a full-thickness placental center cut. This pregnancy only had a single placental disc, an occurrence that happens in approximately 20% of rhesus macaque pregnancies [31] [32]. ISH evaluation of the placental center-cut showed ZIKV RNA in two large villi, one near the trophoblastic shell (Figs 6A and 6B) and the other near the chorionic plate (Fig 6C-F). In both villi, the staining is located predominantly in the villous mesenchyme, and not in the outer syncytiotrophoblast (STB) layer that is directly exposed to maternal blood.

**Figure 6.**
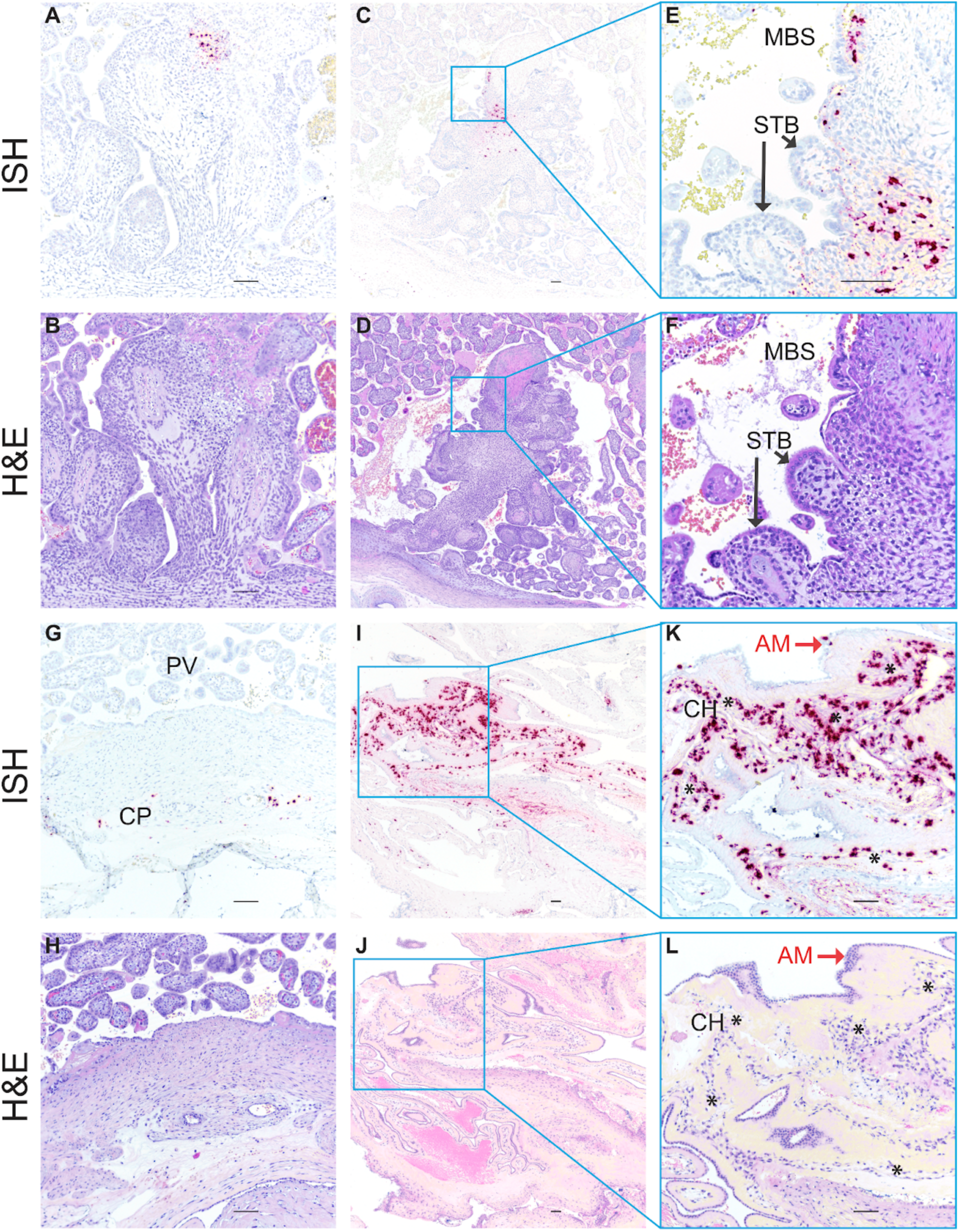
ZIKV RNA detected in MFI by *in situ* hybridization (ISH). Photomicrographs of paraffin embedded sections of the placenta and fetal membranes from 45/14-4. Panels A, C, E, G I, and K Positive ISH indicated by the red/pink chromogenic stain. Panels B, D, F, H, J, and L are corresponding H&E-stained sections to provide histological detail. Panels E, F K and J are higher magnification images of the corresponding regions in the adjacent photomicrographs. Scale bar represents 100 μm. Abbreviations: MBS maternal blood space, STB syncytiotrophoblasts, CP chorionic plate, PV placental villi, AM amnion, CH chorion. Black arrows indicate STB layer of placental villi in Panels E and F, red arrows indicate epithelial cells in the AM in Panels K and L, and * indicate the chorion layer of the fetal membranes in Panels K and L.

Immunohistochemical staining of serial sections of this specimen suggested that ZIKV RNA in the villous near the trophoblastic shell (Fig 6A) may be associated with either macrophages or trophoblasts (S6A Fig, S6B Fig), while the ZIKV RNA in the large villous near the chorionic plate (Fig 6E) localized with cytokeratin-positive trophoblasts (S6E Fig). Additionally, ZIKV RNA was detected sporadically in the chorionic plate surrounding large fetal blood vessels (Figs 6G and 6H).

For 45/14-4, we obtained a high-quality sample of the fetal membranes. The fetal membranes consist of two components: the outer chorion, which is continuous with the chorionic plate of the placenta, and the amnion, the inner membrane that is closest to the fetus and is in contact with amniotic fluid (Fig 4A). While ZIKV was not detected by RT-qPCR, ISH revealed the abundant presence of ZIKV RNA in the fetal membranes (Fig 6 I-L). By comparing the ISH section to an H&E-stained serial section, we see that the ZIKV RNA is mainly detected in the chorion of the fetal membranes, with relatively fewer areas where ZIKV RNA is detected in the amnion (Fig 6 I-L). We confirmed that the ZIKV RNA in the fetal membranes represented replicating virus using mFISH that detects ZIKV replicative intermediates (S5 Fig). Immunohistochemical staining for the macrophage marker CD163 suggests that at least some of the cells with ZIKV RNA may be macrophages (S7 Fig). The relatively abundant presence of ZIKV RNA within the chorion of the fetal membranes and the chorionic plate suggests that ZIKV may gain access to the fetus through the fetal membranes.

### ZIKV-DAK infection correlates with decidual vasculitis and necrosis

To determine the effect of ZIKV-DAK on the development of the fetus, we conducted histological evaluations of the placenta and the fetus and compared our findings to gestational-age-matched controls. Histological evaluation of the placenta was performed by a board-certified pathologist who specializes in human placental pathology; this pathologist was only informed of the gestational age and remained blinded to all ZIKV RNA results and pregnancy treatment. Overall, there was a pattern of increased decidual vasculitis and decidual necrosis in ZIKV cases compared to controls (Table 1). Decidual necrosis was seen in one out of the seven control cases and in four out the nine ZIKV cases. Leukocytoclastic vasculitis in decidual spiral arteries in the decidua was seen in three out of the nine ZIKV cases but not in any of the control cases. We found that decidual vasculitis correlated with the ZIKV RNA within decidual vessels. IHC staining for endothelial cell marker CD31 suggests that the ZIKV RNA is within the endothelial cells in these vessels (S8 Fig). We also found that necrosis of anchoring villi in case 45/14-2 correlated with ZIKV infected macrophages (S9 Fig).

**Table 1.**
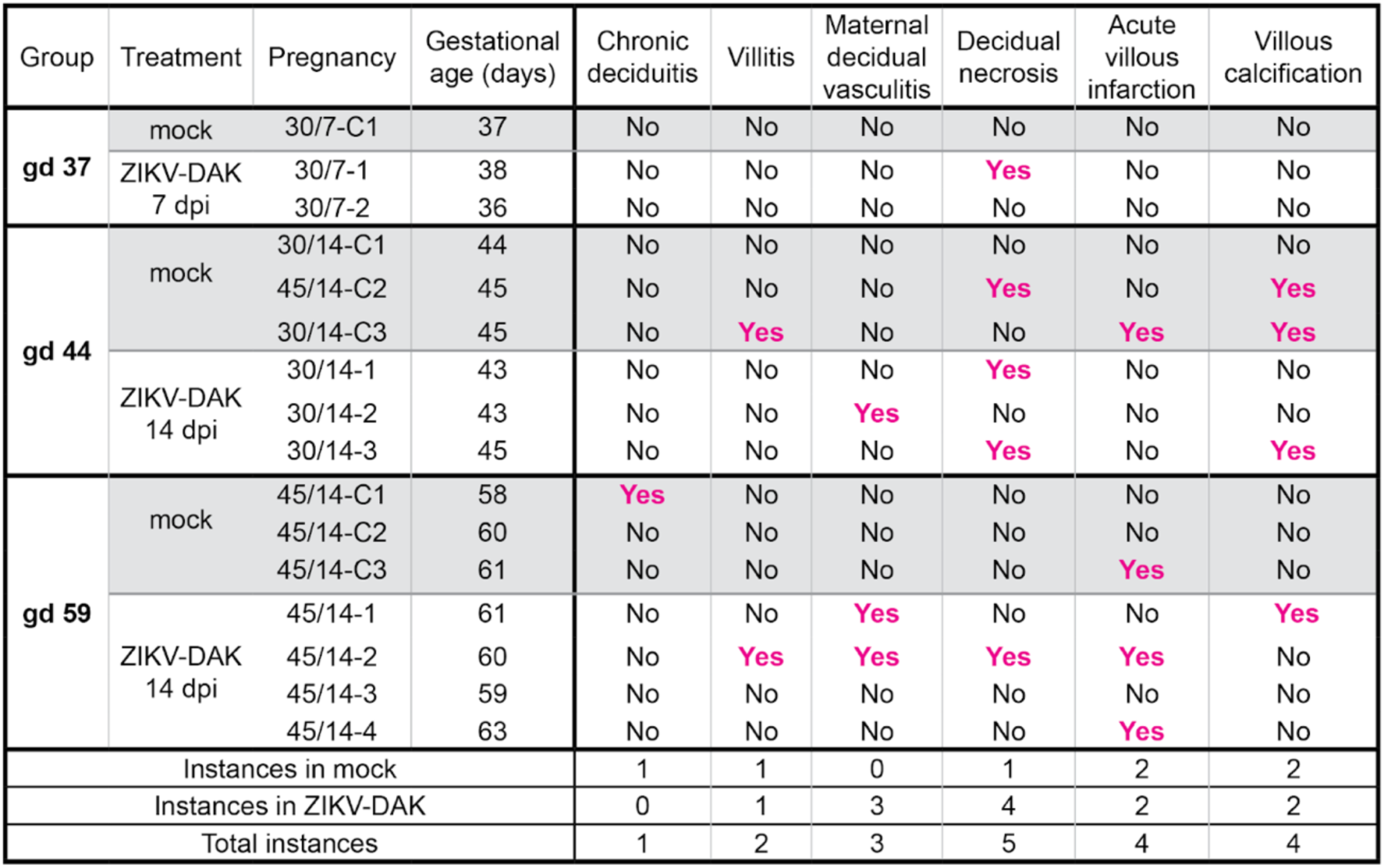
Histopathological evaluation of MFI tissue sections. The presence or absence of specific histological lesions in the MFI are indicated for mock-infected control pregnancies, and ZIKV-inoculated pregnancies. The criteria for villitis excluded inflammation in the anchoring villi, the villi attached to the decidua.

Importantly, we also found ZIKV RNA within decidual spiral arteries in case 30/7-2 (Fig 7). To determine what cell types ZIKV RNA was detected in, we performed IHC to identify extravillous trophoblasts (EVTs), endothelial cells, and macrophages (Fig 7E). IHC staining for CD56 revealed that endovascular EVTs were among the cell types that appeared to be infected with ZIKV. This finding is important because endovascular EVTs play an essential role in spiral artery remodeling [33] [34] and as seen in case 30/7-2 EVTs form “plugs” in early pregnancy that impact uteroplacental blood flow [35].

**Figure 7.**
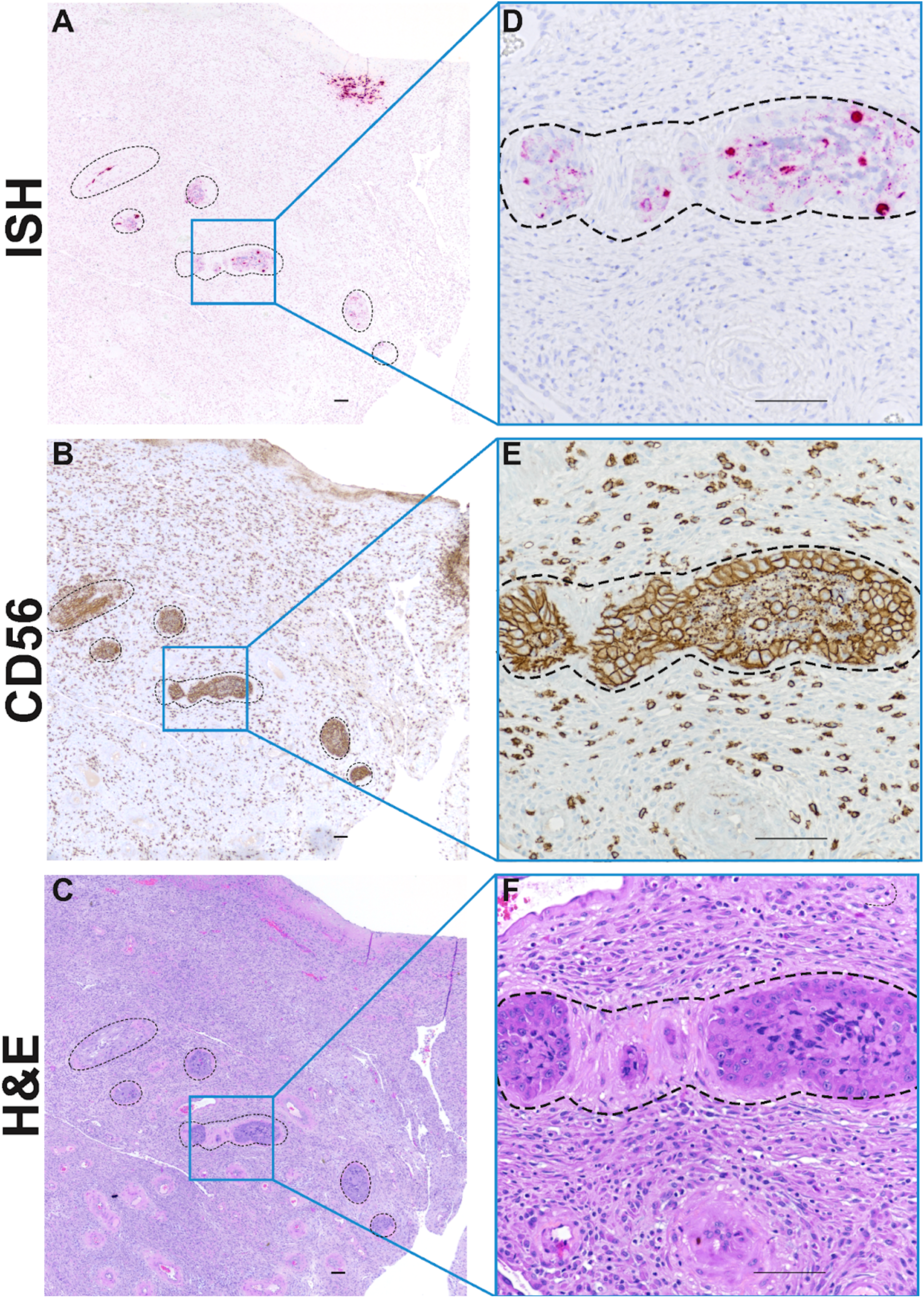
ZIKV RNA in decidual spiral arteries. Photomicrographs of paraffin embedded sections of the decidua basalis from 30/7-2. Spiral arteries are outlined by dashed lines. A) Pink staining shows ZIKV RNA detected using ISH in multiple profiles of a decidual spiral artery with invading luminal extravillous trophoblasts (EVTS). B) Immunohistochemical staining for CD56 reveals large EVT cells as well as smaller immune cells. C) H&E stained serial section. Panels D, E and F represent higher magnification of images of indicated regions of Panels A, B, and C respectively. Scale bar represents 100 μm.

Histological evaluation of the fetus was performed by board-certified veterinary pathologists who were blinded to all RNA results. Eight out of the nine fetuses had no significant histologic lesions when compared to controls. The fetus with multiple ZIKV positive tissues, 45/14-4, had evidence of mild acute hemorrhage in the lung, but this lesion was presumed to be perimortem. Overall, there was no significant pathology in any of the ZIKV exposed fetuses. However, because all nine fetuses were terminated within 14 days of maternal inoculation, we are unable to evaluate what the effects of ZIKV infection in the fetal compartment might have been if the pregnancies had continued.

## Discussion

In this study, we inoculated nine pregnant rhesus macaques with 44 PFU of ZIKV-DAK early in pregnancy at either gd 30 or gd 45 with surgical pregnancy termination at 7 or 14 dpi. All dams became productively infected and developed strong neutralizing antibody responses. Evaluation of the MFI revealed ZIKV RNA in MFI tissues in all pregnancies. Three pregnancies also had infectious ZIKV in the amniotic fluid. One case had ZIKV RNA detected in multiple tissues in the fetus proper. ISH evaluation of MFI tissues showed that the decidua basalis was frequently infected and that the fetal membranes may play a role in ZIKV vertical transmission. No significant histologic pathology in the fetuses was found; however, we found an increase in decidual vasculitis and necrosis in ZIKV-exposed pregnancies compared to gestational-age-matched controls. This study demonstrates that African-lineage ZIKV, like Asian-lineage ZIKV, can be vertically transmitted to the macaque fetus during pregnancy.

### Low-dose inoculation

Productive infection in all dams with inoculation of 44 PFU of ZIKV-DAK suggests maternal infection can occur with a relatively low inoculation dose in macaques. Furthermore, it demonstrates that maternal inoculation with a low dose can achieve vertical transmission, and that vertical transmission with African-lineage ZIKV is not simply a function of a higher dose inoculum as suggested by the comparison of two previously published studies[19] [22]. Vertical transmission was seen in the Raasch et al. study with an inoculation dose of 1X10^8^ PFU [22], but not seen in the Crooks et al. study with an inoculation dose of 1X10^4^ PFU. Styer et al. determined that West Nile virus-infected mosquitoes, when feeding on a live host, inoculate with a dose ranging from as low as approximately 5 PFU to as high as 10^6.6^ PFU, with a median dose ranging from 10^3.4^–10 ^6.1^ PFU depending on mosquito species [36]. Hence, 44 PFU is within the possible range of doses administered by a bite from an infected mosquito, but is likely far from a representative median dose. Although the 44-PFU dose in the current study was unintentional, it has provided novel insight into permissiveness to ZIKV infection in pregnant macaques and risk factors for vertical transmission.

When comparing our data to a study by Crooks et al. [19] in which pregnant rhesus macaques were inoculated with 10^4^ PFU of ZIKV-DAK, we found that although our dams had a statistically significant delay (2.5 days) in the timing of peak viremia, viral RNA reached similar levels in the blood [19]. Additionally, the neutralizing antibody responses observed in the present study were similar to those reported by Crooks et al. [19]. These data suggest that infection with approximately 200 times less infectious virus did not drastically alter the maternal infection dynamics. However, we cannot determine the long-term impact of the inoculation dose on the fetus since pregnancies in this study were terminated within 14 dpi.

### Vertical transmission

The current study suggests that this African-lineage ZIKV isolate, like Asian-lineage ZIKV, can be vertically transmitted to the fetus during pregnancy. ZIKV was vertically transmitted in three pregnancies, as evident by infectious virus in the amniotic fluid. ZIKV RNA was also found in the fetus in two of those three pregnancies. Although the current study used a very small inoculation dose, our results are consistent with what has been found in two other non-human primate studies that show maternal infection with African-lineage ZIKV results in MFI infection and vertical transmission to the fetus [19][20].

Two potential routes of ZIKV vertical transmission have been proposed by Tabata et al.: either through the placenta or through the fetal membranes [37]. These investigators proposed that ZIKV in maternal blood or decidua basalis could reach the placental villi, directly infect the STB layer of the placental villi or the cytotrophoblasts that invade into the decidua to ultimately reach fetal blood vessels within the villous mesenchyme [37]. Alternatively, ZIKV from maternal blood could infect the decidua and spread to the adjacent chorion, the outer layer of the fetal membranes and reach the fetus either through the amniotic membranes and amniotic fluid or by spreading to fetal blood vessels in the chorionic plate in the placenta [37]. In our most apparent case of ZIKV vertical transmission to the fetus (45/14-4), we detected infectious virus in the amniotic fluid and ZIKV RNA in multiple fetal tissues. Focal presence of ZIKV was observed in placental villi, whereas widespread infection was observed in the fetal membranes. This case was the only one where we observed ZIKV in the placental villi by ISH. The morphological evaluation suggests that ZIKV RNA is not present in the outer STB layer but may be in villous macrophages and other cytotrophoblast cells. This pattern of ZIKV in the placenta fits the pattern reported in other *in vivo* studies that found ZIKV infection in the chorionic plate and in the placental villous mesenchyme, but not in the STB layer [19] [38][20]. Importantly, ZIKV RNA was found in a large portion of the fetal membranes in this case, specifically within the chorion. These data suggest that the fetal membranes may act as a ZIKV reservoir and potentially as a route for vertical transmission; however, the broad significance of this observation remains unclear as we did not detect ZIKV RNA in the fetal membranes of the other two cases with infectious ZIKV in the amniotic fluid. We note that only selected biopsies of any tissues were available for analysis in this study, and we suggest that histological approaches that allow for a broader evaluation of the presence of viral genomes in the MFI are important to develop.

### Effect of ZIKV-DAK on maternal-fetal interface histopathology

Studies in mice suggest that congenital infection with African-lineage ZIKV is more likely to cause fetal demise while Asian-lineage is more likely to cause congenital defects[17] [18]. We did not observe any cases of fetal demise, nor did we see any significant pathology in any of the fetuses. However, our study design (low-dose inoculum and early pregnancy termination) may not be ideal for studying adverse fetal outcomes such as fetal loss. Furthermore, the lack of fetal pathology hindered our ability to discern whether maternal inoculation at gd 30 versus gd 45 differentially affected the fetus.

Despite the lack of fetal pathology, we provided evidence that a low dose of ZIKV-DAK does cause pathology in the MFI. We found an increase in decidual vasculitis and necrosis in ZIKV-DAK-exposed pregnancies compared to gestational-age-matched controls. These findings are consistent with those of Hirsch et al. who found decidual vasculitis and necrosis in the MFI in rhesus macaques infected with Asian-lineage ZIKV [39]. Additionally, the potential for ZIKV-DAK to cause pathology in the MFI is illustrated in the cases in which we found ZIKV RNA within trophoblasts located within the lumina of maternal decidual spiral arteries. As far as we are aware, this is the first observation of ZIKV infection of endovascular trophoblasts *in situ* in decidual spiral arteries. Newman et al. reported persistent muscularization in spiral arteries in the decidua of a near-term rhesus macaque pregnancy vaginally challenged with ZIKV-DAK [20]. The importance of the remodeling of these vessels for the establishment of a successful pregnancy [40] [41] provides potential insight into the risk of adverse pregnancy outcomes with early pregnancy maternal ZIKV infection [7] [9].

Congenital birth defects or adverse pregnancy outcomes have not been formally associated with African-lineage ZIKV infection during pregnancy in people. However, African-lineage ZIKV is still considered and monitored as an emerging threat to public health [42]. Like Asian-lineage ZIKV, adverse pregnancy outcomes in people may only be detected when African strains of ZIKV emerge in a ZIKV-naive population. Recently, African-lineage has been found in Brazil, highlighting the risk of this virus to emerge outside the African continent [14]; [15]; [16]. African-lineage ZIKV has been shown to cause fetal demise in *in vivo* studies done in mice and macaques [18] [20], further supporting the potential of African-lineage ZIKV to cause adverse pregnancy outcomes in humans. The current study further supports those findings.

In conclusion, this study demonstrates that African-lineage ZIKV, like Asian-lineage ZIKV, can be vertically transmitted from mother to fetus in the macaque model,, a model that displays key features of human ZIKV infection during pregnancy[24] [28]. The low inoculum dose used in this study demonstrates that the fetal infection observed in previous macaque pregnancy studies using African-lineage ZIKV was not simply the result of administering a large dose [20][19]. Additionally, the low-dose inoculation used in this study suggests a low minimum infectious dose for rhesus macaques and may indicate a high epidemic potential of African ZIKV strains. Finally, using a low-dose inoculation may have the benefit of revealing the most sensitive or susceptible pathway(s) for vertical transmission. It will be instructive to compare the localization of ZIKV at the MFI and in the fetus in these acute low-dose inoculations with early time point collections in pregnant rhesus monkeys receiving higher doses of ZIKV. Understanding how ZIKV compromises the “placental fortress” [43] will help map the pathway to vertical transmission of ZIKV. Understanding the pathway of ZIKV vertical transmission may better prepare researchers and clinicians for future viral outbreaks that may pose a risk for the health of the mother and developing fetus.

## Acknowledgments

We thank the WNPRC Veterinary, Scientific Protocol Implementation, and Pathology Service Unit staff for assistance with animal procedures, including breeding, monitoring, surgery, and necropsy. We thank the University of Wisconsin Translational Research Initiatives in Pathology (TRIP) Laboratory, supported by the UW Department of Pathology and Laboratory Medicine, UWCCC (P30 CA014520) and the Office of Research Infrastructure Programs- NIH (S10 OD023526) for use of its facilities and services. Logan Keding created the illustration of the rhesus macaque conceptus shown in Fig 4A. This research was supported by NIH grants T32 GM007133 to M.R.K., R01 AI132519 to T.G.G, R01 AI132563 to M.T.A and T.C.F, and P51 OD011106 to the Wisconsin National Primate Research Center. The content is solely the responsibility of the authors and does not necessarily represent the official views of the NIH.

## Author Contributions

Conceptualization: Michelle R. Koenig, Emma L. Mohr, Thaddeus G. Golos

Formal analysis: Michelle R. Koenig

Funding acquisition: Emma L. Mohr, Thaddeus G. Golos

Investigation: Michelle R. Koenig, Ann M. Mitzey, Terry K. Morgan, Xiankun Zeng, Heather A. Simmons, Andres Mejia, Fernanda Leyva Jaimes, Chelsea M. Crooks, Andrea M. Weiler

Project administration: Ann M. Mitzey

Resources: Matthew T. Aliota, Thomas C. Friedrich, Thaddeus G. Golos Supervision: Thomas C. Friedrich, Thaddeus G. Golos

Visualization: Michelle R. Koenig, Xiankun Zeng, Logan T. Keding Writing – Original Draft Preparation: Michelle R. Koenig

Writing – Review & Editing: Michelle R. Koenig, Ann M. Mitzey, Terry K. Morgan, Xiankun Zeng, Heather A. Simmons, Andres Mejia, Fernanda Leyva Jaimes, Logan T. Keding, Chelsea M. Crooks, Andrea M. Weiler, Ellie K. Bohm, Matthew T. Aliota, Thomas C. Friedrich, Emma L. Mohr, Thaddeus G. Golos

## Supporting Information Captions

**S1 table. Summary of the timing of inoculation, and the days post infection (dpi) and gestational age at the time of pregnancy termination for each experimental subject.**

**S1 Fig. Timing and magnitude of peak viremia in dams inoculated with 44 PFU or 10^4^ PFU.** A) Timing of peak viremia as measured by RT-qPCR of RNA isolated from plasma. Comparisons were made between dams inoculated with 44 PFU (n=9) and dams inoculated with 10^4^ PFU (n=4) (unpaired t-test, asterisk denotes p = 0.0150). B) Viral load on the day of peak viremia in dams inoculated with 44 PFU (n=9) versus 10^4^ PFU (n=4). The difference was not significant (unpaired t-test, P-value = 0.3729) The horizontal lines within the scatter plots represent the mean for each respective group. 10^4^ PFU data is from Crooks et al.[19].

**S2 Fig 2. Low levels of pre-existing immunity in 45/14-1.** Plaque reduction neutralization tests (PRNT) were performed on serum samples from all nine animals collected prior to ZIKV inoculation. Data are expressed relative to infectivity in the absence of serum. Only positive data is shown.

**S3 Fig. Comparison of neutralizing antibody titers in pregnant rhesus macaques inoculated with 44 PFU vs 10^4^ PFU.** A) PRNT90 values as measured in sera taken approximately 28 dpi from dams inoculated with 44 PFU (n=7) versus 10^4^ PFU (n=4). PRNT90 titers were not significantly different (unpaired t-test, P value = 0.1543). B) PRNT50 values as measured from serum taken approximately 28 dpi from dams inoculated with 44 PFU (n=7) versus 10^4^ PFU (n=4). PRNT50 titers were not significantly different (unpaired t-test, P value = 0.1253). The horizontal lines within the scatter plots represent the mean for each respective group. 10^4^ PFU data is from Crooks et al. [19].

**S4 Fig. Quantification of ZIKV RNA in MFI and fetal tissues.** A) Viral loads determined by RT-qPCR of RNA isolated from tissue in the MFI. B) ZIKV RNA burden in all maternal-fetal interface (MFI) tissues from each group. Tissues include: decidua basalis, uterus-placental bed, placental biopsies, chorionic plate, and fetal membranes. The mean of each group is represented by the horizontal line. C) Viral loads determined by RT-qPCR of RNA isolated from umbilical cord and cord blood.

S5 Fig. ZIKV replicative intermediates detected in fetal membranes from 45/14-4 using multiplex fluorescence *in situ* hybridization (mFISH). Panels A, C, and E show different locations of the slide where ZIKV positive sense RNA (green) and negative sense RNA (red) were detected using mFISH. Nuclei of cells are stained with DAPI (blue). Panels B, D, and F show the ZIKV negative sense RNA detected alone in the same areas. Scale bars in A and E represent 20 μm and the scale bar in C represents 50 μm.

**S6 Fig. Immunohistochemical staining of ZIKV-positive placental villi from 45/14-4.** Photomicrographs of paraffin embedded sections of the placenta from 45/14-4 (please refer to Fig 6A for ISH image). Cytokeratin staining in A), C) and E) identified trophoblasts cells. CD68 staining in B), D) and F) identify placental villous macrophages (Hoffbauer cells). E) and F) show a higher magnification of the corresponding regions of B) and D). Scale bars represent 100 μm.

**S7 Fig. Macrophages in the fetal membranes.** A) photomicrographs of paraffin embedded sections of the fetal membranes from 45/14-4. A) Pink staining shows ZIKV RNA detected using ISH. B) Brown chromogen staining shows macrophages detected using immunohistochemistry staining for CD163. C) H&E stained serial section. Scale bars represent 100 μm.

**S8 Fig. Decidual vasculitis and ZIKV RNA in the decidua.** Decidual vasculitis and ZIKV RNA in the decidua. Representative photomicrographs of paraffin embedded sections of the decidua with vasculitis from 30/14-2. A) and C) Pink staining shows ZIKV RNA detected using ISH in vessels associated with vasculitis. B) and D) H&E stained serial section. Lymphocytic infiltrate is indicated by asterisks. Scale bar represents 100 μm.

**S9 Fig**. Necrosis in anchoring villi and ZIKV RNA in the trophoblastic shell. Representative photomicrographs of paraffin embedded sections of the trophoblastic shell with necrosis in the anchoring villi from 45/14-2. A) Pink staining shows ZIKV RNA detected using ISH in the trophoblastic shell. B) H&E stained serial section. Scale bar represents 100 μm.

## Notes

### Competing Interest Statement

The authors have declared no competing interest.

